# FLASH Irradiation Regulates IFN-β induction by mtDNA via Cytochrome c Leakage

**DOI:** 10.1101/2024.04.10.588811

**Authors:** Jianfeng Lv, Jianhan Sun, Yunbin Luo, Juntao Liu, Di Wu, Yiyu Fang, Haoyang Lan, Longfei Diao, Yuqi Ma, Yuan Li, Meizhi Wang, Ziming Zhao, Heming Wang, Austin Morris, Wenkang Zhang, Zihao Zhang, Lin Lin, Haoyan Jia, Chao Wang, Tianyi Li, Gerard Mourou, Senlin Huang, Gen Yang, Xueqing Yan

## Abstract

Ultrahigh dose rate radiotherapy (FLASH-RT) is under intensive investigation for its biological benefits. The mechanisms underlying its ability to spare normal tissues while suppress tumor growth still remain controversial. Here we reveal that compared to the low dose rate electron irradiation (0.36 Gy/s), FLASH electron irradiation at 61 or 610 Gy/s enhances the cytochrome c leakage from mitochondria in human breast cells MCF-10A, which elicits substantial caspase activation, suppresses both the cytosolic mitochondrial DNA (mtDNA) accumulation and IFN-β secretion. Besides, the deletion of mtDNA severely decreases the radiation-induced cGAS-STING activation. Conversely, the cytochrome c leakage in carcinoma cells MDA-MB-231 post electron irradiation is limited, especially for the case of FLASH irradiation, resulting in less cytosolic cytochrome c but stronger cGAS-STING activation than those in MCF-10A cells. The enhanced difference of cytochrome c leakage between cancer cells and normal cells post FLASH irradiation indicates a potential mechanism of FLASH effect by regulating the apoptotic and inflammatory pathway.

---

Ultrahigh dose rate radiotherapy, also known as FLASH-RT, is a cutting-edge technology in the field of radiotherapy that has garnered extensive global attention during the last decade. Unlike the conventional radiotherapy (CONV-RT) that delivers dose at a relatively low dose rate (typically <0.03 Gy/s, ∼ min), FLASH-RT uses the dose rates several orders of magnitude higher (typically >40 Gy/s, < 0.5 s). The key advantage of FLASH-RT is its ability to yield less damage to normal tissues while maintain equivalent inhibition in tumor growth *in vivo*, which is called as the FLASH effect^1-5^. Clinical trials of FLASH-RT have been launched to treat cutaneous lymphoma by using electrons^6^ and extremity bone metastases by using protons^7^, for its great significance in expanding the therapeutic window by reducing the toxicity inflicted on peripheral tissues. However, for over half a century since the observation of *in vitro* sparing effect by using ultrahigh dose rate irradiation^8^, the physicochemical and biological mechanisms underlying this sparing effect are still a subject of debate within the community^9-11^.

Several hypotheses for the FLASH effect have been proposed from both simulation and experimental work, such as the radiolytic oxygen depletion^9,10^, circulating immune cells protection^12^, and relatively intact DNA integrity post FLASH irradiation^13^. A recent *in vitro* study^14^ discovered that FLASH proton irradiation at 100 Gy/s maintained the morphology and function of mitochondria within normal human lung fibroblasts compared to the case post 0.33 Gy/s irradiation, which could be associated with the radiation induced dephosphorylation of the p-Drp1 protein and mitochondrial fission. Being the site of cellular aerobic respiration and oxidative phosphorylation, mitochondria are the primary source of reactive oxygen species (ROS) within the cells^15^, and they are involved in cell proliferation, death, metabolism and inflammation^16^. Ionizing radiation causes oxidative stress in mitochondria^17^, resulting in mitochondrial outer membrane permeabilization through the opening of BAX or BAK channel^18,19^ and mitochondrial permeability transition pore (MPTP)^20^. This process allows for the cytochrome c and mitochondrial DNA (mtDNA) leakage into cytoplasm, provoking the intrinsic apoptosis via caspase cascade^21^ and type-I interferon (IFN-I) related inflammation via the cGAS-STING pathway^22,23^, respectively. By combining with the apoptotic protease activating factor-1 (APAF-1), cytosolic cytochrome c allows the apoptosome formation and caspase-9 activation, which cleaves the downstream effector caspases and then various cellular substrates. The cyclic GMP-AMP synthase (cGAS), a crucial sensor of cytosol dsDNA, catalyzes the formation of cyclic GMP-AMP (cGAMP) to bind with the STING protein, a stimulator of interferon genes. Phosphorylated STING further interacts with TANK-Binding Kinase 1 (TBK1) and Interferon Regulatory Factor 3 (IRF3), thereby inducing IFN-I expression.

Previous literatures have underscored that radiation-induced cytosolic mtDNA effectively can elicit the anti-tumor immune response^24,25^ and even promote the abscopal effect^26,27^, while some cancer cells have the ability to inhibit the immunogenic effect by enhancing programmed cell death^24,26^. Besides, the suppression of apoptotic caspases enables more type-I interferons production that induced by cytosolic mtDNA^28,29^. Therefore, the opposite functions between cytosolic cytochrome c and mtDNA are crucial to the radiotherapy efficacy for the inflammation regulation. However, it remains unclear how the alteration of radiation dose rates impacts the caspases activation and interferons production in both cancer cells and normal cells. Such an investigation could be important to explore the underlying mechanism of the FLASH effect.

In this study, the direct current superconducting radio frequency (DC-SRF) photocathode gun in Peking University^30^ is utilized to deliver the electron beams with various dose rates. It is discovered that both IFN-β secretion and cytosolic mtDNA accumulation in normal cells MCF-10A are suppressed by FLASH electron irradiation (61-610 Gy/s), which is further mediated by the cytochrome c leakage and caspase activation. In contrast, the results in carcinoma cells MDA-MB-231 post FLASH irradiation are opposite to those in MCF-10A cells, namely, less cytochrome c leakage and more mtDNA accumulation in cytosol. The mechanism of dose rate impact on cytochrome c leakage is further discussed.

## Results

### Irradiation experiment with controllable dose rate and dose delivery

A beam line by using the DC-SRF photocathode gun was designed for electron irradiation experiment (Fig. 1a, Supplementary Fig. 1a). This beam line contains a bending magnet to monitor the daily electron beam energy, which is 1.76±0.03 MeV in this study. Following the bending magnet, a fast-current-transformer (FCT) and a Faraday Cup are applied to monitor the beam’s time structure and integrated current, respectively. The Faraday Cup is retracted after the beam parameters were confirmed, allowing direct beam impingement on a 250 μm thick beryllium window. This window serves a dual role in separating the vacuum environment from the atmosphere and scattering the electrons. After scattering by the beryllium window, the electrons further propagate through a 24 cm aluminum collimator with a 35 mm inner diameter to deliver uniform doses to our samples. Sealed 6-well plates housing the adherent cells are affixed to a 2-dimentional motorized translation stage. Radiochromic films (RCF) are positioned in front of the plates to monitor the dose delivery. The cell layer and RCF are primarily separated by the plate base comprising 1.21 mm thickness of polystyrene, and the corresponding linear energy transfer of electrons in cell layer is 0.186 keV/μm (LET=0.186 keV/μm). Fig. 1b presents the dose distribution in RCF and cells obtained from Monte Carlo simulation with Geant4. The averaged doses in RCF and cells are 0.068 Gy/nC and 0.085 Gy/nC, respectively, resulting in a dose ratio of *D*_cell_ /*D*_RCF_ = 1.25. Examples of RCF dose measurements post irradiation with five beam currents from 4.25 nA to 425 μA are illustrated in Fig. 1c, where the preset dose is 11.5 Gy. The ratio of dose standard deviation and averaged dose for each shot is less than 5%. The beam current can be continuously regulated by mainly changing the repetition rate and charge per micro-pulse (Supplementary Fig. 1b, c). By dividing the measured doses by the corresponding delivery time, the averaged dose rate in RCF exhibits a direct proportionality with the beam current (Fig. 1d), and the linear fitting coefficient is 0.070 Gy/nC, which closely approximates the 0.068 Gy/nC given by Monte Carlo simulations (Fig. 1b). The dose stability is shown in Supplementary Fig. 1d, e.

**Fig. 1.**
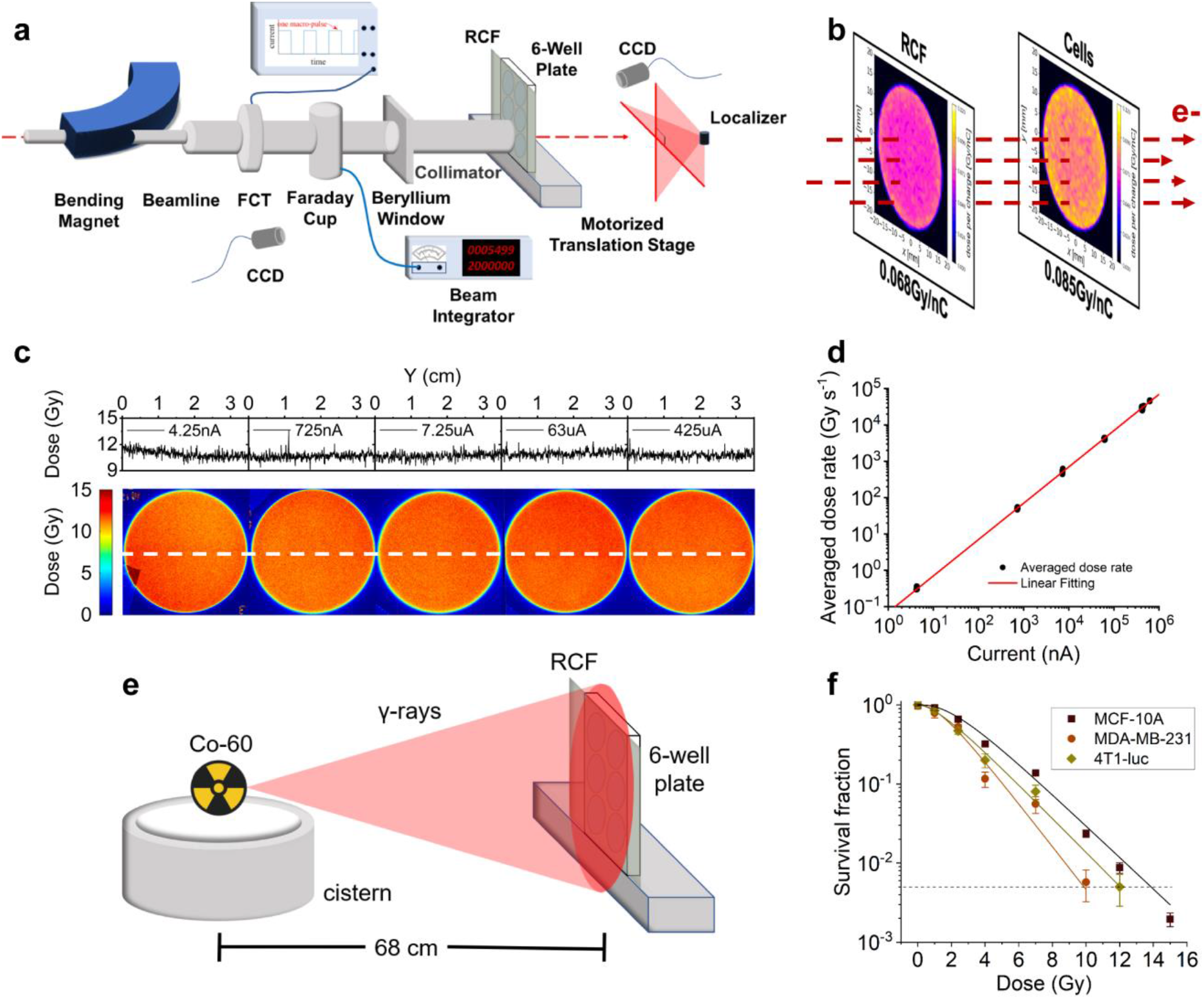
Irradiation experiment with controllable dose rate and dose delivery. **a** Schematic diagram of the electron irradiation setup at the DC-SRF photocathode gun. **b** Monte-Carlo (MC) simulation of the delivered doses in RCF and cell layer. **c** Measured doses in the RCF that post electron irradiation with different beam currents (4.25 nA-425 μA) when the preset dose is 11.5 Gy. **d** The proportional relationship between beam current and the averaged dose rate. **e** Schematic diagram of the Cobalt-60 γ-rays irradiation. **f** Clonogenic assays of three cell lines by using Cobalt-60 γ-rays irradiation, fitting curves are given by the single-hit, multi-target model model. (n=3 biologically independent samples). Data in **f** are presented as mean±SD.

The Cobalt-60 γ-rays irradiation experiment (Fig. 1e) at the Cobalt Laboratory of Peking University was performed to investigate the radio-sensitivity of the cell lines used in this study (LET=0.2 keV/μm, 0.36 Gy/s), including the non-tumorigenic human breast epithelial cells MCF-10A, human breast carcinoma cells MDA-MB-231, and tumorigenic mouse mammary carcinoma cells 4T1-luc with luciferase expression. Clonogenic survival fractions of MCF-10A, 4T1-luc, and MDA-MB-231 cells present radio-sensitivity differences and result in an equivalent survival fraction (5‰) at delivered doses of 14 Gy, 12 Gy, and 10 Gy, respectively (Fig. 1f). These dose data are used in the electron irradiation experiment to further evaluate the consequential effects of varying dose rates. Fitting curves of the survival fraction in Fig.1e are obtained by using the single-hit, multi-target model^31^ SF=1-(1-exp(-*kD*))^*n*^, the fitting parameters (*k, n*) for MCF-10A, MDA-MB-231, and 4T1-luc cell lines are (0.46, 2.94), (0.63, 2.43), and (0.49, 1.91), respectively.

### Interferon-β production depends on the electron irradiation dose rate

The cGAS-STING activation post electron irradiation was investigated to find out the potential dose rate impact, as reported by Shi et al. in their FLASH X-ray experiments^13^. The fluorescence intensity of the cytosolic dsDNA extract was normalized to that of the corresponding whole-cell extract under irradiation with different dose rates. However, the cytosolic dsDNA levels within neither MCF-10A nor MDA-MB-231 cells exhibited significant changes post irradiation with different dose rates (Fig. 2a). Intriguingly, despite exposure to radiation doses resulting in an equivalent survival fraction (5‰), MDA-MB-231 cells demonstrated a much larger amount of cytosolic dsDNA accumulation compared to MCF-10A cells (Fig. 2b). Specifically, the cytosolic dsDNA fraction in MDA-MB-231 cells almost exceeded 50% post 10 Gy irradiation, whereas after 14 Gy irradiation, the detected cytosolic dsDNA in MCF-10A cells was generally lower than 40%. The phosphorylated STING protein (pSTING) within the cells was then observed and quantified via immunofluorescence imaging (Fig. 2c, d). As to the MCF-10A cells 9 hours after electron irradiation, the cells of 0.36 Gy/s group has the largest fluorescent pSTING area compared to the cells of FLASH group (61 Gy/s and 610 Gy/s), while such a difference cannot be observed for the MDA-MB-231 cells (Fig. 2e). Furthermore, the IFN-β concentration in the cell culture supernatant was measured 48 hours after the electron irradiation. FLASH irradiation lead to reduced IFN-β secretion in MCF-10A cells compared to the 0.36Gy/s irradiation. In MDA-MB-231 cells, however, the supernatant IFN-β concentration exhibits uptrends with the increase of irradiation dose rate.

**Fig. 2.**
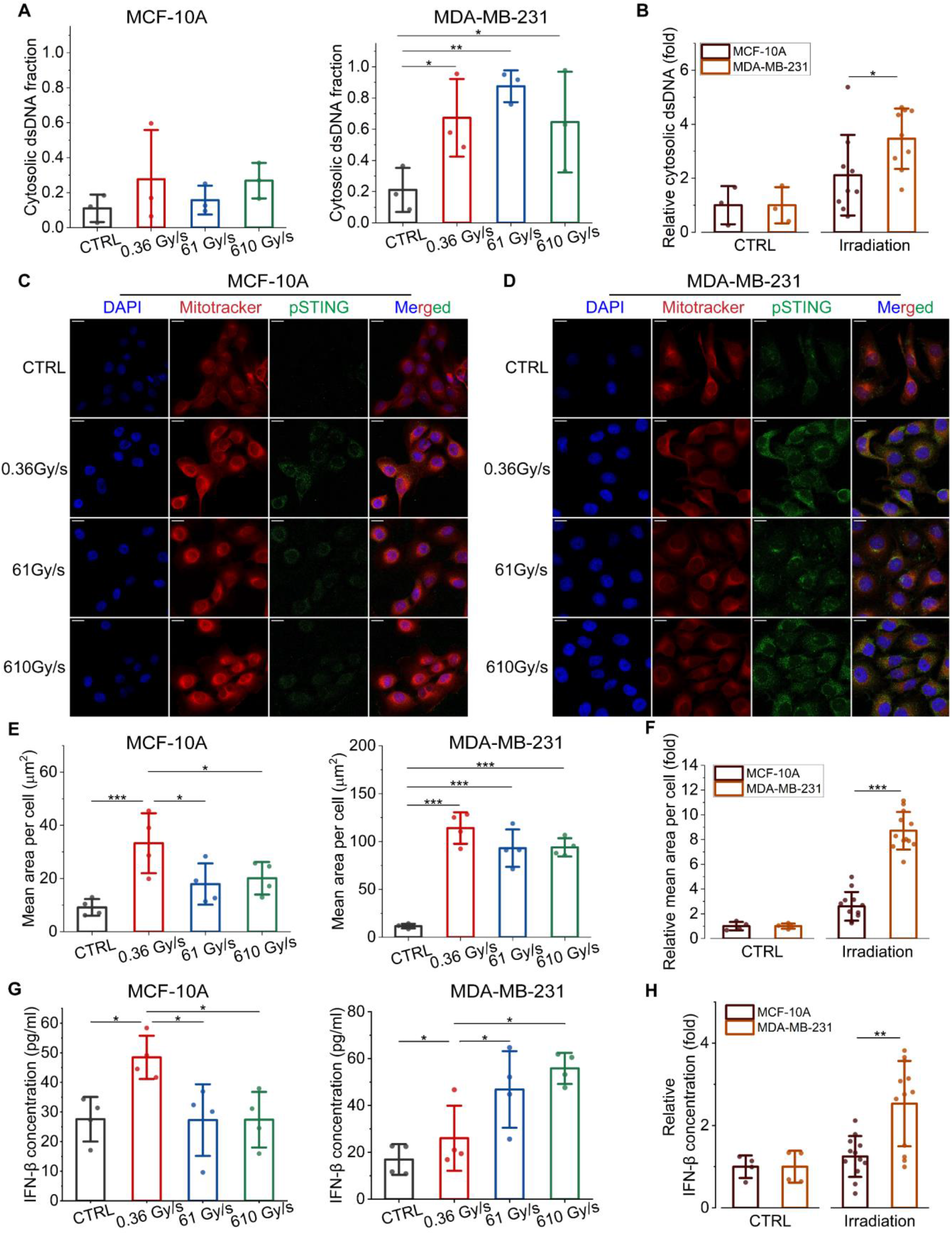
Electron irradiation activates the cGAS-STING pathway. **a** The cytosolic dsDNA accumulation of MCF-10A cells (14 Gy) and MDA-MB-231 cells (10 Gy) 48 hours after irradiation with indicated dose rates (n=3 biologically independent samples). **b** Comparison of cytosolic dsDNA fraction between MCF-10A cells and MDA-MB-231 cells as shown in **a**. (**c, d**) Representative immunofluorescence images (blue: nuclei, red: mitochondria, green: p-STING) of MCF-10A and MDA-MB-231 cells 9 hours after irradiation with indicated dose rates. Scale bars: 20 μm. **e** Quantification of p-STING fluorescence area per cell as shown in **c, d** (n=4 biologically independent samples). **f** Comparison of relative p-STING fluorescence area between MCF-10A cells and MDA-MB-231 cells as shown in **e. g** Supernatant IFN-β concentration of MCF-10A and MDA-MB-231 cells 48 hours post irradiation with indicated dose rates. (n=4 biologically independent samples). **h** Comparison of relative IFN-β concentration between MCF-10A cells and MDA-MB-231 cells as shown in **g**. Statistical significance was calculated via unpaired t-test in **b, f, h**, and one-way ANOVA test in **a, e, g**. Data in **a, b, e, f, g, h** are presented as mean±SD. *p<0.05, **p<0.01, ***p<0.001.

It is noted that the carcinoma cells MDA-MB-231 presented stronger cGAS-STING activation compared to the normal cells MCF-10A post electron irradiation, for both the relative p-STING fluorescence area and IFN-β secretion of MDA-MB-231 significantly surpass those of MCF-10A cells (Fig. 2f, h), which are consistent with the observation of stronger cytosolic dsDNA accumulation in MDA-MB-231 cells (Fig. 2b). These phenomena indicate a dose rate dependent mechanism impacting the IFN-β production, and even leading to the different response in cGAS-STING activation between MCF-10A cells and MDA-MB-231 cells.

### Electron irradiation dose rate affects the cytosolic mtDNA accumulations and Interferon-β production through apoptotic caspases

The dose rate dependent IFN-β production was found to be inconsistent with the cytosolic dsDNA fraction in Fig. 2a. To explore the origination of the cytosolic dsDNA, specific primers targeting nuDNA (β-actin, GAPDH) and mtDNA (ND1, D310) were used in real time quantitative PCR, and the cytosolic dsDNA fraction was normalized against the non-irradiated control group. For MCF-10A cells, cytosolic mtDNA accumulation decreases significantly in the 61 Gy/s irradiation group compared to the 0.36 Gy/s irradiation group (Fig. 3a). Conversely, for MDA-MB-231 cells, irradiation with an ultrahigh dose rate of 61 Gy/s increases mtDNA accumulation in cytosol (Fig. 3b). However, such a difference is not significant in the nuDNA accumulation. The nuclear damage was further investigated by detecting the phosphorylated form of histone H2AX (γH2AX) protein (Supplementary Fig. 3), which serves as a sensitive marker to nuclear DNA double strand breaks. All the nuclei suffered strong damage post irradiation, and γH2AX fluorescence appeared in a large fraction of each nucleus. Additionally, statistical analysis shows no noticeable difference in the γH2AX area per nucleus post irradiation with different dose rates.

**Fig. 3.**
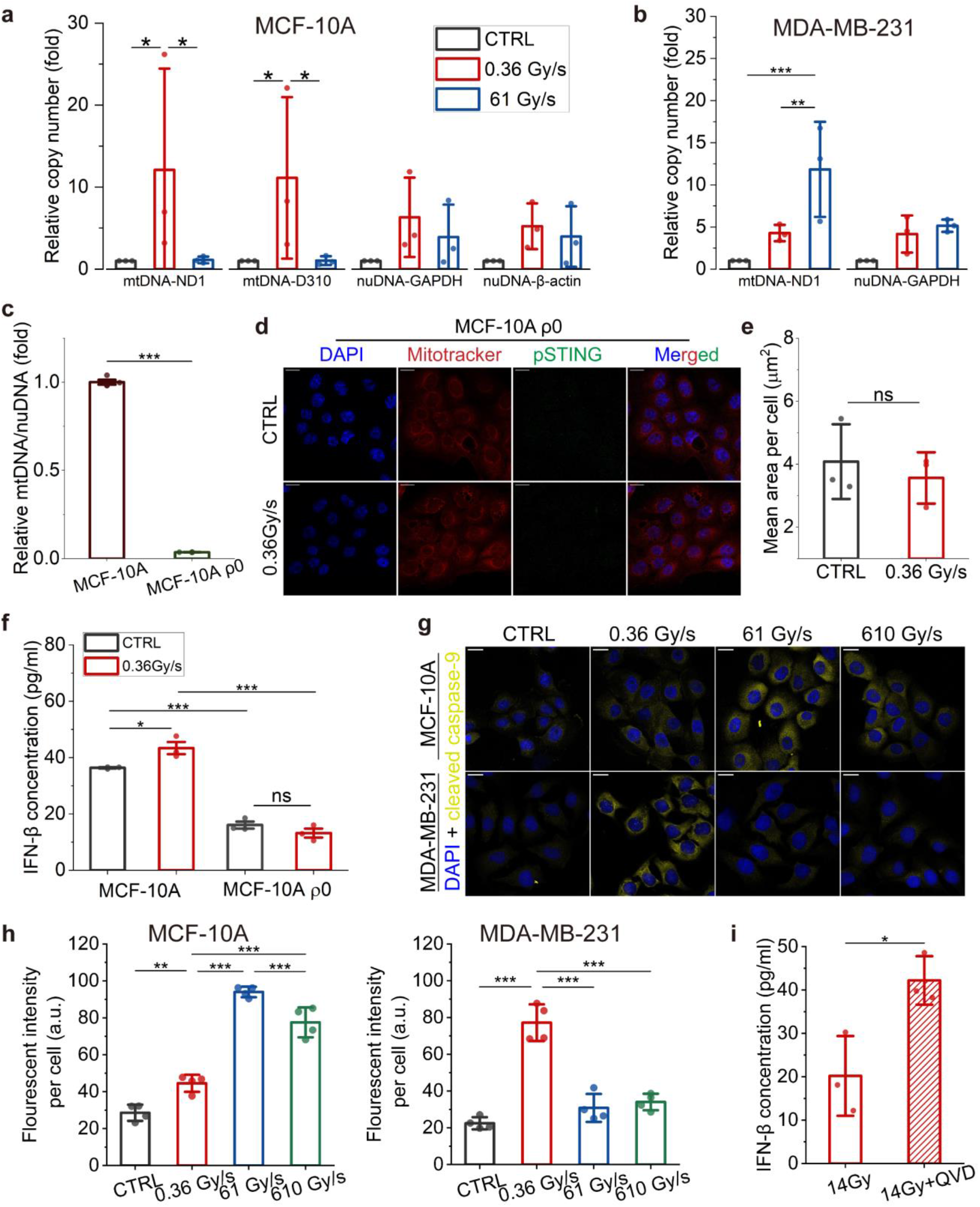
Electron irradiation induces caspases activation and cytosolic mtDNA accumulation. Cytosolic fraction of nuclear dsDNA and mitochondrial dsDNA for **a** MCF-10A cells and **b** MDA-MB-231 cells 48 hours after electron irradiation with indicated dose rates (n=3 biologically independent samples). **c** Comparison of the mitochondrial dsDNA level between MCF-10A cells and MCF-10A ρ0 cells. **d** Representative immunofluorescence images (blue: nuclei, red: mitochondria, green: p-STING) of MCF-10A ρ0 cells 9 hours after 0.36 Gy/s irradiation. Scale bars: 20 μm. **e** Quantification of p-STING fluorescence area per cell as shown in **d** (n=3 biologically independent samples). **f** Supernatant IFN-β concentration of MCF-10A and MCF-10A ρ0 cells 48 hours after 0.36 Gy/s irradiation. (n=3 biologically independent samples). **g** Representative immunofluorescence images (blue: nuclei, yellow: cleaved caspase-9) of MCF-10A and MDA-MB-231 cells 9 hours after electron irradiation with indicated dose rates. Scale bars: 20 μm. **h** Mean fluorescent intensity of cleaved caspase-9 per cell as shown in **g**. (n=4 biologically independent samples). **i** Supernatant IFN-β concentration of MCF-10A cells 48 hours after 0.36 Gy/s irradiation with 14 Gy and treatment with caspase inhibitor QVD-OPh. Statistical significance was calculated via unpaired t-test in **c, e, i**, and one-way ANOVA test in **a, b, f, h**. Data in **a, b, c, e, f, h, i** are presented as mean±SD. *p<0.05, **p<0.01, ***p<0.001.

The change in mtDNA accumulation, likely attributable to the mitochondrial response to electron irradiation, emphasizes the significance of mitochondria and potentially serves as a mechanism of FLASH effect by reducing type-I interferon related inflammation in normal tissues. The MCF-10A ρ0 cells, characterized by the mtDNA absence, were also utilized for comparative investigation to show the mtDNA importance in radiation-induced IFN-β production. qPCR analysis revealed a significant decrease in mtDNA level in MCF-10A ρ0 compared to the normal MCF-10A cells (Fig. 3c). The irradiated and non-irradiated MCF-10A ρ0 cells have no difference in both the STING phosphorylation and IFN-β production, while the supernatant IFN-β concentration of MCF-10A cells exhibits a significant increase after 0.36 Gy/s irradiation (Fig. 3f and Fig. 2g). Moreover, the supernatant IFN-β concentration of MCF-10A ρ0 cells is much lower than the normal MCF-10A cells, which indicates the deficiency of cGAS-STING pathway without mtDNA.

Additionally, these changes of mtDNA accumulation with irradiation dose rate are found to be accompanied by the opposite change of apoptotic caspases activation. Specifically, compared to the 0.36 Gy/s irradiation, FLASH irradiation results in more cleaved caspase-9 in MCF-10A cells, but significantly mitigates the caspase-9 activation in MDA-MB-231 cell (Fig. 3g, h). The apoptotic caspases has been extensively proved to enable the type-I interferons suppression^24,28,29,32^ and even dsDNA degradation^33,34^ in previous literatures. To confirm the caspase importance in this study, the complete culture medium containing caspases inhibitor (QVD-OPh) was added to the adherent MCF-10A cells immediately after 0.36 Gy/s irradiation and then cultured for 48 hours. The irradiated cells with caspase inhibition released much more IFN-β to the supernatant culture medium (Fig. 3i) than the cells with irradiation only. Through enhancing the caspases activation, FLASH irradiation suppresses the mtDNA accumulation and IFN-β production in MCF-10A cells.

### Cytochrome c leakage is enhanced in MCF-10A but suppressed in MDA-MB-231 by FLASH electron irradiation

The cytochrome c leakage from mitochondria is an important inducer for caspases activation and intrinsic apoptosis. To investigate the influence of irradiation dose rate on cytochrome c leakage, the immunofluorescence of cytochrome c and complex V, an inner mitochondria membrane protein that won’t release during apoptosis, was applied 9 hours post electron irradiation. Cells without irradiation show strong colocalization between cytochrome c and complex Vα subunit, while the cells post electron irradiation present significant cytochrome c leakage to cytosol and mitochondrial networks enlargement (Fig. 4a, f). This morphological change is due to the mitochondrial fission and mass increase under stress^17^. The Manders’ colocalization coefficient (MCC), which is the ratio of colocalized pixels intensity in green channel and all pixels intensity in green channel, as well as the relative cytosolic cytochrome c were calculated for totally 40 fields of each group. Compared to 0.36 Gy/s irradiation, FLASH irradiation results in smaller MCC and stronger cytochrome c leakage in MCF-10A (Fig. 4b, c). Besides, the released cytochrome c proteins in FLASH groups pervasively distribute throughout the entire cytoplasm (Fig. 4a). In contrast, opposite phenomena occur in MDA-MB-231 cells. Compared to 0.36 Gy/s irradiation, the FLASH irradiation reduces the cytochrome c leakage significantly. The MCC are maintained high post FLASH irradiation, and the pervasive distribution of cytochrome c appealed in MCF-10A cells can be hardly observed in MDA-MB-231 cells. These results in both MCF-10A and MDA-MB-231 cells showed strong consistency among the cytochrome c leakage and caspase-9 activation.

**Fig. 4.**
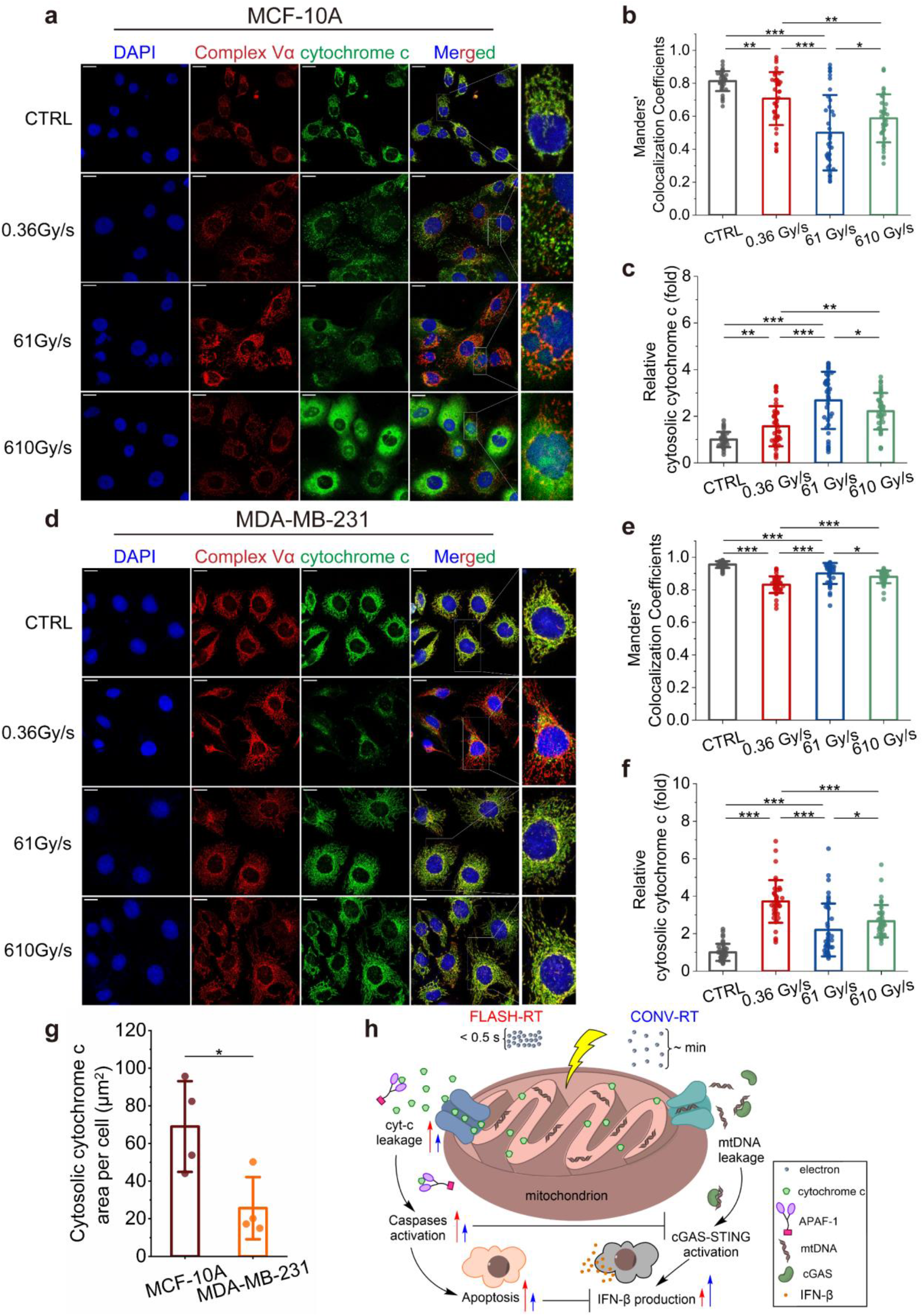
Cytochrome c leakage is regulated by electron irradiation dose rate. **a** Representative immunofluorescence images (blue: nuclei, red: complex Vα, green: cytochrome c) of MCF-10A cells 9 hours after electron irradiation with indicated dose rates. Scale bars: 20 μm. (**b, c**) Colocalization analysis of complex Vα subunit and cytochrome c as shown in **a**. (n=4 biologically independent samples, 10 fields for each sample). **d** Representative immunofluorescence images (blue: nuclei, red: complex Vα, green: cytochrome c) of MDA-MB-231 cells 9 hours after electron irradiation with indicated dose rates. Scale bars: 20 μm. (**e, f**) Colocalization analysis of complex Vα and cytochrome c as shown in **f** (n=4 biologically independent samples, 10 fields for each sample). **g** Comparison of cytosolic cytochrome c between MDA-MB-231 and MCF-10A cells after 0.36 Gy/s irradiation as shown in **a, d**. (n=4 biologically independent samples. **h** The mechanism of reduced IFN-β production in MCF-10A cells post FLASH irradiation by enhancing the cytochrome c leakage. Statistical significance was calculated via unpaired t-test in **g**, and one-way ANOVA test in **b, c, e, f**. Data in **b, c, e, f, g** are presented as mean±SD. *p<0.05, **p<0.01, ***p<0.001.

The mitochondrial morphology change after irradiation was also analyzed from the complex Vα fluorescence by using ImageJ^35^ (Supplementary Fig. 3). Two types of mitochondrial structures, including individuals and networks, were considered. Individuals include the large, round, rod and punctate mitochondria, and networks are the structures with at least one junction and three branches. No significance of the morphology change with irradiation dose rate was found in the MCF-10A cell. As to MDA-MB-231 cells, both the number of mitochondrial individuals and networks per cell increased with the irradiation dose rate, and the mean branches per network were fewer in the FLASH group, which indicated the enhanced mitochondrial fission and reduced morphology network complexity post FLASH irradiation. Even so, the 0.36 Gy/s irradiation, not FLASH irradiation, leads to more cytochrome c leakage in MDA-MB-231 cells (Fig. 4f), while the cytosolic area of cytochrome c is still significantly smaller than that in MCF-10A cells (Fig. 4g). This difference in cytochrome c leakage could be related to the Warburg effect^36^ that cancer cells tendentiously produce energy by aerobic glycolysis in cytoplasm rather than oxidative phosphorylation and citric acid cycle in mitochondria. Besides, cancer cells are usually more resistant to mitochondrial membrane permeabilization inducers that lead to cytochrome c leakage^37^. The different mitochondrial response to ionizing radiations between normal cells and cancer cells could be pivotal in the FLASH effect.

As for the MCF-10A cells in the study, the FLASH irradiation leads to the increase of cytochrome c leakage and the subsequent caspases activation (Fig. 4h), which then suppresses the cGAS-STING pathway and IFN-β production. Interestingly, the results in MDA-MB-231 post FLASH irradiation are totally the opposite, namely, less cytochrome c leakage and caspases activation, which allow the enhanced cGAS-STING activation. The potential physicochemical mechanism underlying the dose rate dependent cytochrome c leakage is further discussed in the following section.

## Discussion

The occurrence of FLASH effect entangles two dimensions consisting of dose delivery method and physiological function of different cells. It is noted that in quantitative assessment including all cytosolic dsDNA, no change of the cytosolic dsDNA was found with the increase of radiation dose rate for both MCF-10A and MDA-MB-231 cells (Fig. 2a, b). Instead, a subtle investigation using qPCR revealed the different cytosolic mtDNA accumulation post irradiation with different dose rates (Fig. 3a, b), since mitochondrial DNA accounts for only 0.25% of the whole human cellular DNA^38^. In the prior investigations conducted by Shi et al.^13^ involving FLASH X-rays irradiation (120 Gy/s), a significant reduction in cytosolic dsDNA of intestinal organoids was observed through PicoGreen fluorescence quantification, which is a much stronger indicator than what we observed. Nevertheless, the normal human breast cell line MCF-10A used for our study is immortalized rather than the primary cells isolated from animal tissues, such that genes involved in cell cycle regulation, apoptosis, or DNA repair may exhibit elevated expression levels, and increase the intrinsic genome instability. Additionally, The oxygen partial pressure at the irradiation site is considered to be important in the FLASH effect^39-41^, due to their roles in the indirect damage to cellular DNA^42^. A hypoxic environment is usually needed to result in less nuclear DNA damage^43-46^. the oxygen-rich environment of 2D cultures in our experiment places the cells in an enhanced state of radio-sensitivity, making it difficult for nuclear DNA damage to be influenced by the irradiation dose rate.

Irradiated tumor cells have the ability to suppress intrinsic DNA sensing by hijacking caspase 9 signaling^24^, while our results in carcinoma cells MDA-MB-231 show that FLASH electron irradiation can better reduce the caspase-9 activation and then increase the IFN-β production compared to the low dose rate irradiation. Interestingly, there are also evidences about the improved tumor control by using FLASH protons irradiation^47,48^. We further investigate this potential by including a vaccination model in mice (Supplementary Fig. 4). 3 million irradiated 4T1-luc cells were subcutaneously injected into the left flank of each mouse as the vaccine, and the control group cells were sham irradiated. Seven days after the vaccination, one million untreated 4T1-luc cells were injected into the right flank of each mouse to rechallenge. Compared to the mice in the control group, the mice in irradiation groups showed delayed tumor formation, and the *in vivo* tumor growth curves have no significance with the irradiation dose rate (Supplementary Fig. 4d). Living imaging with bioluminescence revealed the presence of cancer cells in the right flank (the rechallenge site) of several mice in the control group (Supplementary Fig. 4c). In contrast, the mice in irradiation groups did not demonstrate any tumor development on the right flank over a four-week period, irrespective of the variation in radiation dose rate. This finding supports the equivalent, but not the enhanced immunogenicity of 4T1-luc cancer cells *in vivo* by using FLASH electron irradiation. The anti-tumor efficacy of FLASH-RT might be different in different cancer cell lines, but at least can be equivalent to that of conventional low dose rate radiotherapy.

The importance of cytochrome c and mitochondria dysfunction in ultrahigh dose rate irradiation was firstly demonstrated by Han et al.^49,50^, they found that the loss of cytochrome c significantly reduced late apoptosis and necrosis of mouse embryonic fibroblast cells post the proton irradiation at 10^9^ Gy/s, and the difference was more pronounced than that with 0.05 Gy/s ^60^Co γ-ray irradiation. In another in vitro experiment carried out by Guo et al.^14^, they observed the protection of FLASH proton irradiation (LET = 10 keV/μm, 100 Gy/s) on mitochondrial morphology of normal human diploid lung fibroblasts, which is associated with the reduced dephosphorylation of the phosphorylated Dynamin-1-like protein, while the underlying physicochemical mechanism of the dose rate impact on dephosphorylation process remains unbeknown. In our electron irradiation experiment with dose rate ranging from 0.36 Gy/s to 610 Gy/s, however, no difference in the mitochondrial morphology change was observed in MCF-10A cells, and even enhanced mitochondrial fission was observed in MDA-MB-231 cells post FLASH irradiation. Furthermore, the variation of cytochrome c leakage with increased irradiation dose rate was found to be inconsistent with the mitochondrial morphology change. These phenomena cannot be explained by the hypothesis about p53-Drp1 pathway proposed by Guo et al^14^.

Therefore, we proposed the hypothesis of electron transport chain disruption to elucidate the potential mechanism of the FLASH effect through the radiation induced cytochrome c leakage (Supplementary Fig.5). The cytochrome c leakage from mitochondria proceeds by a two-step process^51^, firstly, the detachment of cytochrome c and cardiolipin (CL) to generate a soluble pool of cytochrome c; secondly, the permeabilization of mitochondrial outer membrane to enable the release of cytochrome c. The electron transfer in cytochrome c oxidase (Complex IV, a key component of the electron transport chain) has a time scale of millisecond or larger^52,53^, which is comparable to the time delivery of FLASH-RT, but much smaller than that of CONV-RT. The normal healthy cells are characterized by strong function of electron transport chain (ETC) in mitochondria^37^. Post the FLASH irradiation, the ETC function could be disrupted immediately due to the water radiolysis induced chemical reaction and diffusion, which occur within a second. In contrast, the ETC function can be maintained to a much longer time during CONV-RT before complete disruption. Consequently, FLASH-RT leads to quick ATP consumption and the inner mitochondrial membrane (IMM) depolarization, therefore the opening of permeability transition pore^20^ to allow the leakage of cytochrome c and mtDNA. Moreover, Excessive ROS production after the ETC dysfunction stimulates the CL peroxidation and cytochrome c detachment in a relative short time, which results in a cascade feedback from ETC dysfunction to cytochrome c leakage, and finally enhances the intrinsic apoptosis but suppresses the type-I interferon related inflammation, thus contributing to the normal tissues sparing effect. As to most of the cancer cells, the aerobic glycolysis in cytosol, but not the oxidative phosphorylation in mitochondria, provides cancer cells with most of the energy supply (Warburg effect^36^). The mitochondrial CL anomalies are accompanied by marked decreases in the activities as well as functional capacities of electron transport chain components^54^. FLASH-RT could still disrupt the ETC function immediately, but neither initialize substantial cytochrome c detachment nor then stimulate a positive feedback, such that the cytochrome c leakage cannot be stronger than that in CONV-RT.

## Methods and Materials

### Cell culture

The non-tumorigenic human breast epithelial cells MCF-10A were kindly gifted from Dr. Mingjie Gao. The cells were grown in DMEM/F12 (Pricella) containing 5% HS, 20ng/mL EGF, 0.5μg/mL Hydrocortisone, 10μg/mL Insulin, 1% non-essential amino acids and 1% penicillin/streptomycin (P/S). To prepare the mtDNA depleted MCF-10A cells, the culture medium was further supplemented with 50 ng/ml ethidium bromide, 1mM sodium pyruvate and 50 μg/ml uridine, and the cells were incubated for 5 passages. Human breast carcinoma cells MDA-MB-231 were cultured in Eagle’s Minimum Essential Medium (DMEM) supplemented with 10% FBS and 1% P/S. The 4T1 cells with luciferase expression (4T1-luc) were used for the tumor model on the mice BALB/c strain. 4T1-luc cells were cultured in RPMI 1640 Medium supplemented with 10% FBS and 1% P/S.

One day before the irradiation experiment, cells were either seeded on the confocal petri dish with a 15-mm glass bottom (NEST, no. 801002) for immunofluorescence imaging or seeded on the 6-well plate for other assays. All the cells were incubated at 37 °C with 5% CO_2_ and 21% O_2_.

### Ultra-high dose rate electron irradiation

The electron irradiation experiments were performed at the Superconducting Radio-frequency (SRF) Accelerator Laboratory of Peking University^30^, where the photocathode gun can produce stable electron beams with the energy of mega-electron-volt (MeV). The repetition rate of electron micro-pulses is determined by the driving laser, which can work at two modes of 1 MHz and 81.5 MHz. The micro-pulse duration (picosecond to sub-picosecond) and charge per micro-pulse (picocoulomb) can also be continuously adjusted by the driving laser. In our experiment, we adopted the setup consists of a scatterer and a collimator to deliver uniform dose, and the maximal averaged dose rate achieved was at the order of 10^4^ Gy/s.

### Dose monitoring and calculation

The electron energy was measured by a bending magnet in the beamline, and the time structure was monitored by a fast-current-transformer and Faraday Cup. Radiochromic films (RCF, EBT3 type) were placed in front of the cell samples for dose measurement. An Epson Perfection V700 scanner was used to scan the irradiated RCF in the transmission mode, and the calibration of RCF dose response to ionization radiations was referenced to the previous literatures^55,56^. Due to the electron energy decrease from RCF to cells, the delivered dose in cells differed from the dose in RCF. The absolute doses absorbed by cells were calculated by Monte Carlo simulations with Geant4^57-59^, in which we constructed the whole electrons transportation from the vacuum to cells.

### Clonogenic cell survival assay

Cobalt-60 γ-ray was used to assess these cell lines’ radio-sensitivity in the culture conditions. The absorbed dose rates at different distances away from the radioactive source were previously calibrated in the Cobalt-60 laboratory of Peking University. Cells for each group were seeded in the 6-wells plates one day before the irradiation. Irradiation groups were irradiated under the dose rate 10 Gy/min, and the non-irradiated groups were blocked with lead bricks. The linear energy transfer (LET) of Cobalt-60 γ-ray in cells is about 0.2 keV/μm. After irradiation, cells were cultured for 10 days and the culture media were replaced with the fresh every 3 days. Then the adherent cells were fixed with 4% paraformaldehyde for 10 minutes and dyed with 0.5% crystal violet. The colonies whose diameters were larger than 0.3 mm were counted to calculate the survival fraction (SF).

The single-hit, multi-target model is formulated as SF=1-(1-exp(-*kD*))^*n*^, where *D* is the absorbed dose, where n is the number of sensitive sites or targets in a cell, *k=*1*/D*_0_ and *D*_0_ is the dose that lead to an average of one hit per site.

### Immunofluorescence

At the indicated time post electron irradiation (1 hour for phosphorylated H2AX (γH2AX) observation, 9 hours for phosphorylated STING, cytochrome c, and cleaved caspase-9 observation), cells were fixed with 4% paraformaldehyde for 10 minutes for storage or immediate use. Immunofluorescence processes were referenced to the antibody product instructions. Briefly, Samples were permeabilized with 0.1% Triton X-100 for 10 minutes at 4 °C, and then were blocked with the 10% normal goat serum for 45 minutes at room temperature before incubating with the primary antibodies (anti-γH2AX: Beyotime, no. C2035S-4; anti-pSTING: Cell Signaling Technology, no. 40818S; anti-cytochrome c and anti-complex V: Abcam, no. Ab110417; anti-cleaved caspase-9: Affinity, no. AF5244) at 4 °C overnight. In the following day, samples were further incubated with corresponding secondary antibodies for 1 hour at room temperature. Finally, the samples were mounted with mounting medium containing DAPI (Beyotime, no. P0131). The fluorescence was detected with an LSM-700 confocal microscope (Zeiss) and analyzed with ImageJ.

### Cytosolic dsDNA quantification

48 hours post irradiation, total DNA and cytosolic DNA were extracted and quantified as previously described^25,60^. Briefly, cells collected from 6-wells plate were divided into two equal aliquots. The first aliquot was suspended in 500 μL of 50 mM NaOH and boiled for 30 min to dissolve the DNA. Then, 50 μL of 1 M tris-HCl (pH 8.0) was added to balance the pH and centrifuged at 17,000 g for 10 min to separate intact cells. These extracts were used to quantify total dsDNA. The second aliquot was suspended in 500 μL of buffer with 150 mM NaCl, 50 mM Hepes (pH 7.4), and digitonin (25 μg/ml; HARVEYBIO, no. LS1463). The mix were then incubated end-over-end for 10 min on ice to enable selective membrane permeabilization and centrifuged at 980 g for 3 minutes 3 times to pellet intact cells. Finally, the cytosolic supernatants were transferred to new tubes and spun at 17,000 g for 10 min to pellet any leftover cellular debris. All dsDNA samples were purified with DNA Clean & Concentrator-5 (ZYMO RESEARCH, no. D4013) before further use. The PicoGreen dsDNA reagent (200× dilution) (Thermo Fisher Scientific, no. P11496) was used for dsDNA quantification with fluorescence. The fluorescence intensity is proportional to the dsDNA concentration in solution and obtained by a multifunctional microplate reader (Thermo Fisher Scientific).

### Quantitative real-time PCR

To detect the possible differences of nuDNA and mtDNA leakage to cytoplasm under different irradiation dose rates, the quantitative real-time PCR (qPCR) was performed on 7500 FAST real-time PCR system with Tag Pro Universal SYBR Master Mix (Vazyme, no. Q712-02) according to the instruction manuals. The qPCR samples were prepared with the same method described in the section of cytosolic dsDNA quantification. Ct values for whole-cell extracts served as normalization controls for the values obtained from the cytosolic extracts. The nuDNA and mtDNA primers used in qPCR are listed below. nuDNA β-actin(F): ACCCACACTGTGCCCATCTAC; nuDNA β-actin(R): TCGGTGAGGATCTTCATGAGGTA; nuDNA GAPDH(F): AGCCACATCGCTCAGACACCA; nuDNA GAPDH(R): GCAAATGAGCCCCAGCCTTC; mtDNA ND1 (F): CACCCAAGAACAGGGTTTGT; mtDNA ND1 (R): TGGCCATGGGTATGTTGTTAA; mtDNA D310 (F): CACAGACATCATAACAAAAAATTTCC; mtDNA D310 (R): GGTGTTAGGGTTCTTTGTTTTTGG.

### Interferon-β detection

After the irradiation, 1 ml fresh culture medium was added to each well containing 2×10^6^ to 3×10^6^ cells. Then the cells were incubated for 48 hours before transferring the media supernatants to fresh tubes. These IFN-β samples were centrifuged at 1000 g for 20 minutes for immediate use or storage at -20 °C. The human IFN-β ELISA kit was used to quantify the IFN-β concentrations within the supernatants. ELISA processes were referenced to the instruction manual.

### Animal experiment

Six-week-old female BALB/c mice were bought from Beijing Vital River Laboratory Animal Technology Co., Ltd. A vaccination model was employed to examine the proliferative capacity and the immunogenic effectiveness of 4T1-luc cells *in vivo*, subsequent to irradiation at varying dose rates. Following an exposure to 12 Gy under three different dose rates, namely, 0.36 Gy/s, 61 Gy/s, and 610 Gy/s, a total of 3×10^6^ 4T1 cells were subcutaneously inoculated to each mouse on the right flank. The 4T1-luc cells for the control group experienced sham radiation. One week post vaccination, the mice were challenged by subcutaneously injecting 10^6^ non-irradiated 4T1-luc cells into the left flank. Tumor growth was recorded every two days over a period of four weeks and terminated at the point of mice euthanasia. Tumor volume was calculated using the formula *V=*0.52*×L×W*^2^, where *L* and *W* are the tumors longest and shortest diameters, respectively. Three weeks post the initial inoculation, bioluminescence living imaging was availed to visualize the distribution of the cancer cells within the mice.

### Ethics statement

All animal experiments and procedures were performed in accordance with “Guiding Principles in the Care and Use of Animals” (China) and approved by the Peking University Institutional Animal Care and Use Committee (Approval ID: Physics-YangG-2).

### Statistical analysis

Origin (version 2022) (OriginLab Corp., Northampton, MA, USA.) was used for statistical analysis and graph generation. Each experiment was repeated with at least three biologically independent samples. Comparison between two groups was performed using unpaired *t*-test. Comparison between outcomes of different dose rates was performed using one-way analysis of variance (ANOVA) with Holm-Bonferroni’s multiple comparisons test. All the quantitative results were presented as mean±SD. For all graphs, *p<0.05; **p<0.01; and ***p<0.001.

## Code availability

The code used to analyze mitochondrial morphology networks has been deposited and reported in the manuscript by Valente et al. The full source code of ImageJ macros for analyzing immunofluorescence images are available on GitHub: https://github.com/Jeffrey-Lv/code-for-FLASH-experiment.

## Data availability

All data generated or analyzed during this study are included in this article. Raw data are available from the corresponding author upon reasonable request.

## Acknowledgement

We acknowledge the funding received from the National Key Research and Development Program of China (No. 2023YFC2413200/2023YFC2413201 and 2019YFF01014400), National Natural Science Foundation of China (No. 12375334 and 11921006), Open Research Fund Project of the State Key Laboratory of Nuclear Physics and Technology, PKU (No. NPT2022ZZ01; NPT2020KFY19 and NPT2020KFJ04).

## Author contribution

J.Lv contributed to the overall project design, performed experiments and data analysis, and wrote the manuscript. G.Y. and X.Y. designed and supervised the overall project, performed data analysis, and edited the manuscript. S.H. designed the electron beam line and supervised the accelerator operation. J.S. and Y.L. performed experiments and data analysis with J.Lv. J.Liu, L.L, H.J. and L.T. contributed to the beam line design and beam control. D.W., H.L. and M.W. performed experiments with J.F.L. F.Y., L.D., Y.M., Z.Zhang, and A.M. contributed to the dose monitoring and analysis. Y.M., Z.Zhao, H.W. and W.Z. contributed to the sample preparation and data analysis. H.W. performed the experiments with interferon-β. Y.L. contributed to the experiments with BALB/c mice. G.M. contributed to the research guidance and discussion.

## Additional information

### Competing interests

The authors declare no competing financial interests related to this manuscript.

**Correspondence and requests** for materials should be addressed to Senlin Huang, Gen Yang or Xueqing Yan.

