## Supplementary Figures for "FLASH Irradiation Regulates IFN-β induction by mtDNA via Cytochrome c Leakage"

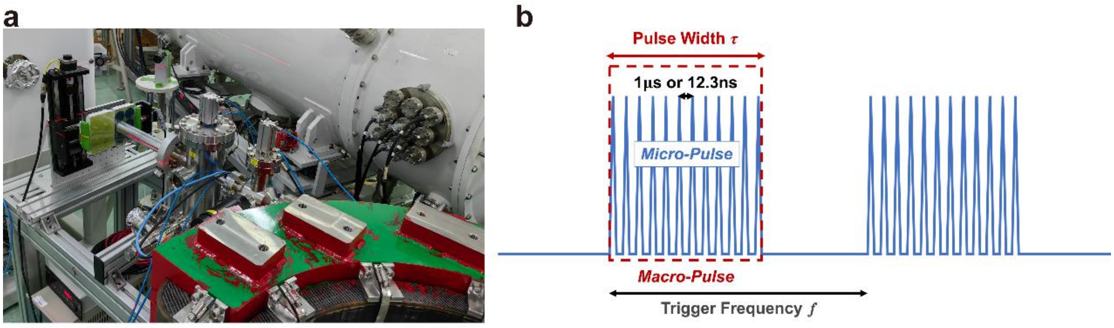


c

| | | | | | |
|---|---|---|---|---|---|
| Pulse width τ (ms) | 0.1 | 240 | 24 | 2.9 | 0.39 |
| Trigger frequency f (Hz) | 10 | 0.1 | 0.1 | 0.1 | 0.1 |
| Repetition rate (MHz) | 1 | 1 | 1 | 81.25 | 81.25 |
| Beam current I (μA) | 0.0042 | 0.72 | 7.2 | 62 | 430 |
| Charge per micro-pulse (pC) | 4.2 | 0.72 | 7.23 | 0.76 | 5.3 |
| Instantaneous dose rate (Gy/s) | $3.6\times10^{7}$ | $6.2\times10^{6}$ | $6.2\times10^{7}$ | $6.5\times10^{6}$ | $4.5\times10^{7}$ |
| Averaged dose rate (Gy/s) | 0.36 | 61 | 610 | 5270 | 36550 |

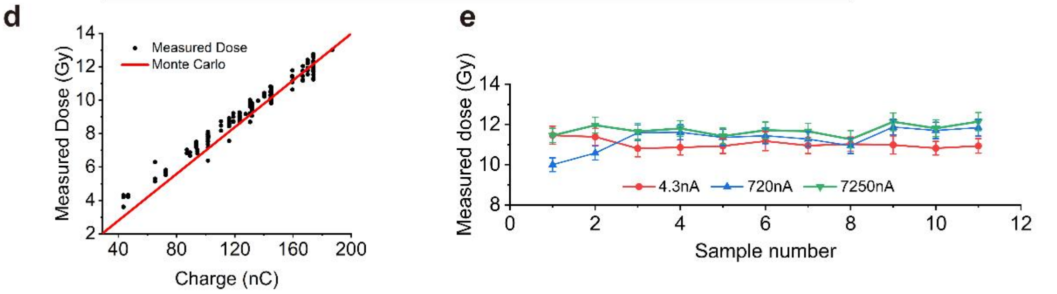


**Supplementary Fig. 1 | Electron beam parameters and dose delivery to radiochromic films. a** The time structure of typical electron beams, which can work at two modes of repetition rate, i.e. 1 MHz and 81.5 MHz (corresponding temporal separations are 1 μs and 12.3 ns). **b** Beam parameters of five delivery strategies spanning 5 orders of magnitude of beam current. **c** The dose stability of 10 consecutive shots when the preset dose is 11.5 Gy. **d** The comparison between dose measurements from RCF and Monte-Carlo simulation.

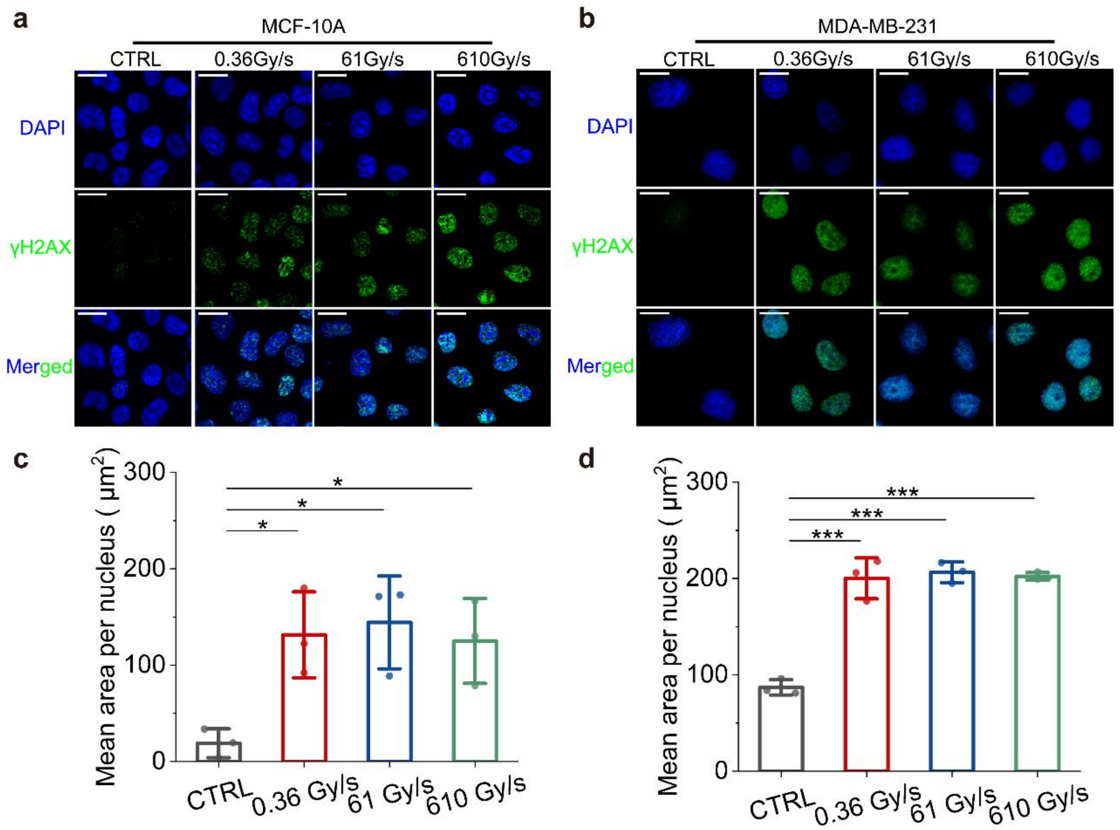


**Supplementary Fig. 2 | Nuclear DNA double strand breaks generated by electron irradiation.** (**a, b**) Representative immunofluorescence images (blue: nuclei, green: γH2AX) of nuclei γH2AX of MCF-10A and MDA-MB-231 cells 1 hours post irradiation with indicated dose rates. Scale bars: 20 μm. (**c, d**) Quantification of γH2AX fluorescence area per cell as shown in **a**, **b** (n=4 biologically independent samples). Statistical significance was calculated via one-way ANOVA test in **c**, **d**. Data in **c**, **d** are presented as mean±SD. *p<0.05, **p<0.01, ***p<0.001.

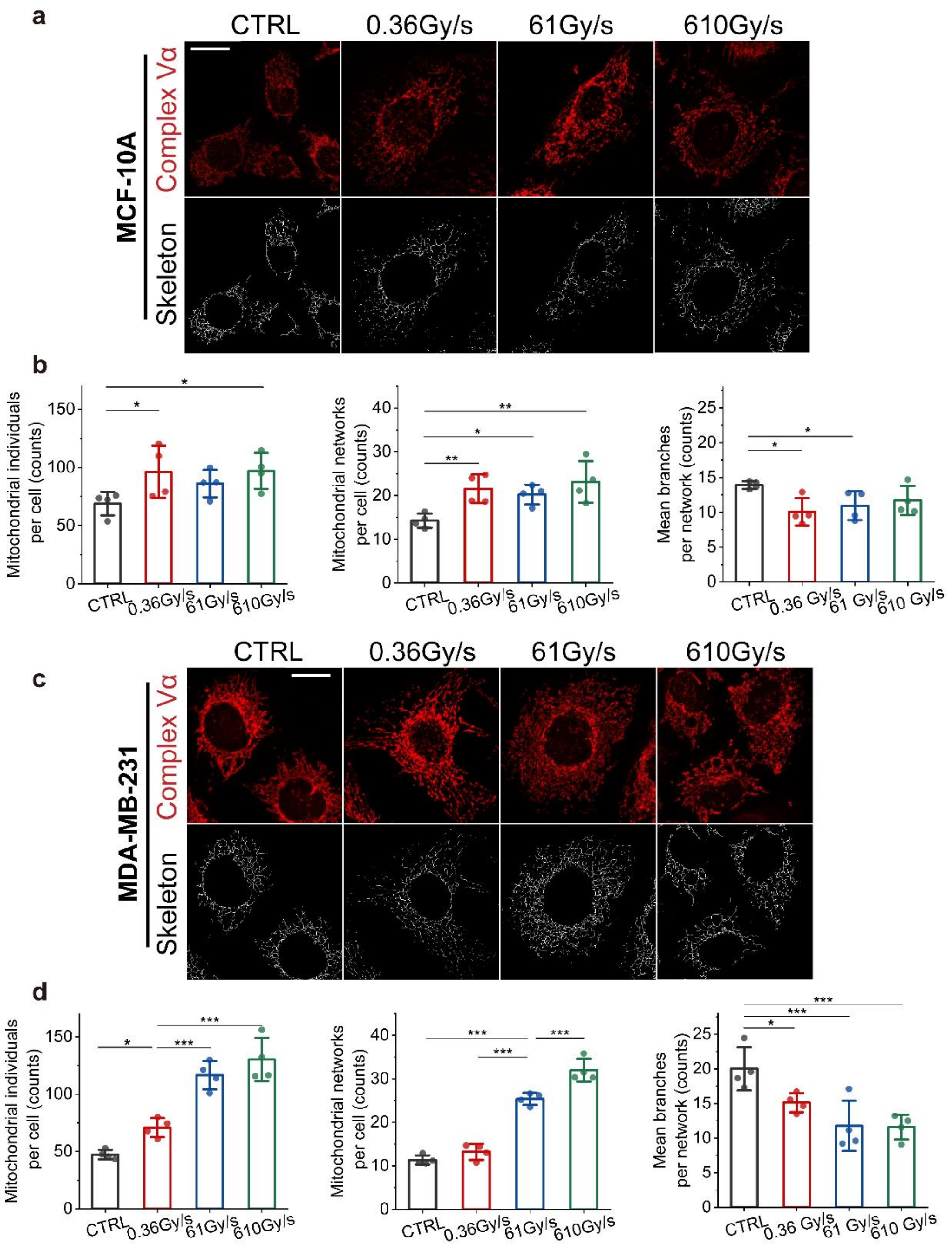


**Supplementary Fig. 3 | Mitochondrial morphology change post electron irradiation. a** Representative immunofluorescence images of MCF-10A mitochondria 9 hours post irradiation (red: mitochondria labelled by complex Vα unit), and the corresponding skeletonized mitochondria images. Scale bars: 20 μm. **b** Statistical analysis of the MCF-10A mitochondria morphology from the skeletonized mitochondria (n=4 biologically independent samples). **c** Representative immunofluorescence images of MDA-MB-231 mitochondria 9 hours post irradiation (red: mitochondria labelled by complex V α unit), and corresponding skeletonized mitochondria images. Scale bars: 20 μm. **d** Statistical analysis of the MDA-MB-231 mitochondria morphology from the skeletonized mitochondria (n=4 biologically independent samples). Statistical significance was calculated via one-way ANOVA test in **b**, **d**. Data in **b**, **d** are presented as mean±SD. *p<0.05, **p<0.01, ***p<0.001.

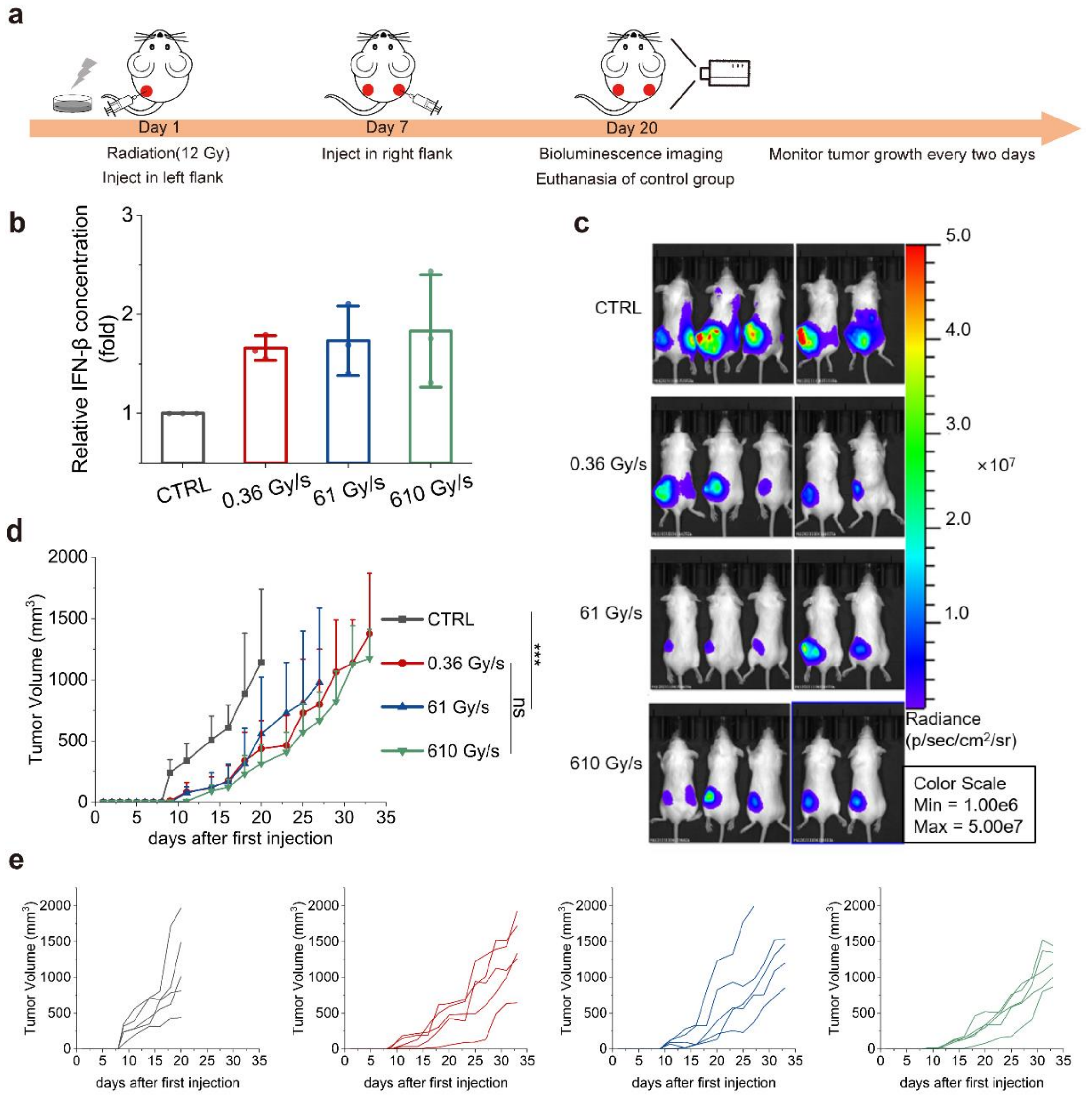


**Supplementary Fig. 4 | The anti-tumor vaccination effect of electron irradiation. a** Scheme for the study of vaccination effect of electron irradiation. **b** Supernatant interferon-β concentration *in vitro* after the irradiation (12 Gy) with indicated dose rate. (n=3 biologically independent samples). **c** Living imaging of the tumors by bioluminescence at Day 20. **d** Averaged tumor growth curves in the left flanks after the irradiation (12 Gy) with indicated dose rate. (n=5 biologically independent animals). **e** Individual tumor growth curves in **d**. Statistical significance was calculated via one-way ANOVA test in **b**, and unpaired linear mixed effect modeling in **d**, and Data in **b**, **d** are presented as mean±SD. *p<0.05, **p<0.01, ***p<0.001.

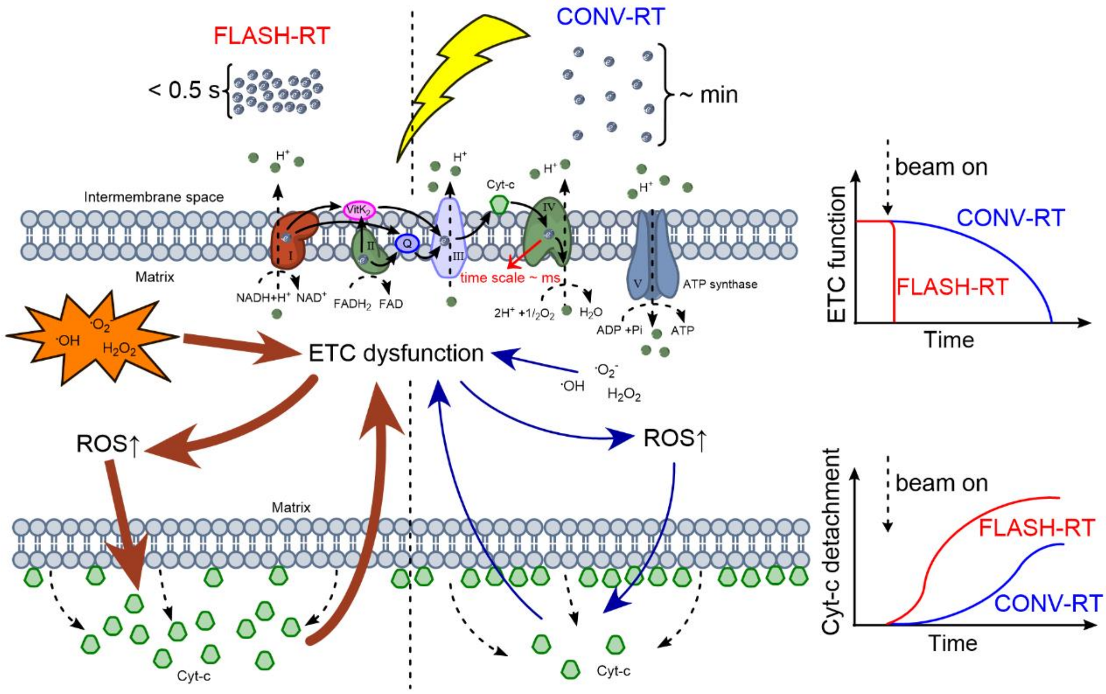


**Supplementary Fig. 5 | The electron transport chain hypothesis in the dose rate dependent cytochrome c leakage.** FLASH-RT stimulates an extensive cascade feedback of the electron transport chain (ETC) dysfunction and cytochrome c detachment from cardiolipin in the normal cells, which enhances the cytochrome c leakage from mitochondria to cytosol. CONV-RT leads to a small quantity of cytochrome c detachment events during the whole irradiation, which limits this cascade feedback.
